# Dietary fatty acids promote lipid droplet diversity through seipin enrichment in an ER subdomain

**DOI:** 10.1101/424663

**Authors:** Zhe Cao, Yan Hao, Yiu Yiu Lee, Pengfei Wang, Xuesong Li, Kang Xie, Wen Jiun Lam, Yifei Qiu, Guanghou Shui, Pingsheng Liu, Jianan Qu, Byung-Ho Kang, Ho Yi Mak

## Abstract

Exogenous metabolites from microbial and dietary origins have profound effects on host metabolism. Here, we report that a sub-population of lipid droplets (LDs), which are conserved organelles for fat storage, is defined by metabolites-driven targeting of the *C. elegans* seipin ortholog, SEIP-1. Loss of SEIP-1 function reduced the size of a subset of LDs while over-expression of SEIP-1 had the opposite effect. Ultrastructural analysis revealed SEIP-1 enrichment in an endoplasmic reticulum (ER) subdomain, which co-purified with LDs. Analyses of *C. elegans* and bacterial genetic mutants indicated a requirement of polyunsaturated fatty acids (PUFAs) and microbial cyclopropane fatty acids (CFAs) for SEIP-1 enrichment, as confirmed by dietary supplementation experiments. In mammalian cells, heterologous expression of SEIP-1 promoted lipid droplet expansion from ER subdomains in a conserved manner. Our results suggest that microbial and polyunsaturated fatty acids serve unexpected roles in regulating cellular fat storage by enforcing LD diversity.

---

Lipid droplets (LDs) are the primary organelles for fat storage in all eukaryotic cells ^1–3^. They accommodate excess energy in the forms of triglycerides (TAG) and cholesterol esters (CE) within a phospholipid monolayer that is enriched in phosphatidylcholine or phosphatidylethanolamine ^4–6^. Extensive proteomic analyses revealed a core set of proteins that reside on the surface of LDs, some of which regulate fat deposition and mobilization ^7^. Although LDs play a central role for intracellular fat storage, they are known to accommodate additional cargos in a developmental stage- and tissue-specific manner ^8–10^. The diversification of LDs is further hinted by the preferential association of metabolic enzymes to subsets of LDs ^11,12^. However, the molecular mechanisms that underlie LD diversity are not fully understood.

In the energy replete state, LDs are closely associated with the endoplasmic reticulum (ER) ^13,14^, which can be facilitated by protein-protein interactions or through stable or transient membrane connections ^12,15,16^. The physical coupling of the two organelles is required for LD expansion, in part because TAG precursors are made in the ER before they are converted into TAG by DGAT2 at the ER-LD interface ^12,15,17,18^. Additional evidence for a fundamental role of the ER in cellular fat storage comes from the discovery that Atlastins regulate LD size ^19^. Atlastins are conserved proteins essential for the generation and maintenance of the tubular ER network ^20^. Depletion of Atlastins reduces LD size ^19^, possibly due to a loss of tubular ER subdomains that are dedicated for ER-LD coupling. However, how such subdomains are maintained and segregated from the rest of the network is unknown.

The seipin protein family plays a conserved role in lipid homeostasis and maintenance of lipid droplet morphology. Mutations in seipin/BSCL2 cause a severe form of congenital generalized lipodystrophy in humans ^21,22^. Pleiotropic metabolic defects such as insulin resistance, hypertriglyceridemia, and hepatic steatosis are thought to stem from an almost complete lack of adipose tissues. Accordingly, seipin has been shown to act cell-autonomously to promote adipocyte differentiation ^23–25^. The human seipin protein was reported to reside in the ER when over-expressed in mammalian cells ^26^. Intriguingly, seipin orthologs were found at ER-LD junctions where it may be responsible for the proper partitioning of lipids and proteins between the ER and LDs ^22,27–30^. In addition, loss of seipin function in yeast and *Drosophila* alters the profile of acyl chains in phospholipids ^28,31^, which has been proposed to cause gross disturbance to LD morphology, resulting in ‘supersized’ LDs or clusters of abnormally small and misshapen LDs ^28,30^. More recently, a role for seipin in the maturation of nascent LDs has been reported ^32^. Therefore, the precise localization of seipin appears to be critical for its function.

The phospholipid composition is a major determinant of the structure and function of eukaryotic membrane-bound organelles ^33^. The synthesis of phospholipids is in turn dependent on the availability of exogenous and endogenous precursors of head groups and fatty acyl chains in the form of fatty acids ^34,35^. In humans, dietary polyunsaturated fatty acids are readily incorporated into membrane phospholipids. They modulate membrane fluidity and serve as precursors for key immunomodulatory molecules. In addition to the diet, the gut microbiome is another potential source of exogenous fatty acids. For example, bacterial cyclopropane fatty acids are found to be incorporated into phospholipids and triglycerides in human adipose tissue ^36^. However, the significance of bacterial fatty acids in eukaryotic membranes is so far obscure.

In this paper, we report that human seipin and its *C. elegans* ortholog SEIP-1 are targeted to a tubular ER subdomain. SEIP-1 is enriched in ER tubules that tightly associate with a subset of LDs. Proper targeting of SEIP-1 is critically dependent on specific polyunsaturated fatty acids (PUFAs) and bacterial cyclopropane fatty acids. Therefore, LDs heterogeneity may originate from fatty acids-driven targeting of distinct protein ensembles.

## Results

### SEIP-1 regulates LD size

A single seipin ortholog, SEIP-1, was identified in the *C. elegans* proteome by BLAST. Membrane topology predictions suggested that, similar to human seipin, SEIP-1 has two transmembrane helices flanking a conserved central loop region, which may reside in the ER lumen (Figure S1A) ^37,38^. To elucidate the function of SEIP-1 in *C. elegans*, we examined *seip-1(tm4221)* mutant animals, which harbored a deletion that removed exon 4 and part of exon 3 of the R01B10.6 open reading frame (Figure 1A). We could not detect full-length SEIP-1 by Western blotting in these animals (Figure 1E). Furthermore, RNA interference (RNAi) against *seip-1* in *seip-1(tm4221)* animals did not modify their phenotypes (Z.C., unpublished data). Therefore, *tm4221* is likely a genetic null allele. We focused our phenotypic analysis on the intestine, which is the major site of fat storage in *C. elegans* ^2^. Using label-free Stimulated Raman Scattering (SRS) ^39^, we did not detect significant difference in neutral fat storage between *seip-1* mutant and wild type animals (Figure 1B). The functionality of the SRS system was validated for its ability to detect elevated fat storage in *daf-22* mutant animals, which are known to store more triglycerides ^40^. We extended our analysis beyond neutral lipids by comparing wild type and *seip-1* mutant animals with a mass spectrometry-based lipidomic approach (Figure S1B). The relative amount of two classes of lipids were significantly increased in *seip-1(-)* animals: phosphatidic acid (PA) and diacylglycerol (DAG) (Figure S1C). To investigate the impact of such perturbation in lipid profile on LD morphology, we used mRuby::DGAT-2 as a marker ^15^. The median size of LDs in the intestinal cells was reduced from 0.81μm in diameter in wild type animals to 0.42μm in *seip-1(-)* animals (Figure 1C-D, 1H). The alteration of LD size in *seip-1(-)* animals were reproducibly observed when we used DHS-3::GFP as an alternative LD marker (Figure S1E-G) ^41^. Taken together, the abundance of small LDs correlated with the over-representation of phospholipids with small head groups (PA and DAG), possibly because they are more compatible with high membrane curvature. Nevertheless, not all LDs were reduced in size in *seip-1(-)* animals (Figure 1H, inset). A similar phenotype of heterogeneous LDs was observed in seipin deficient budding yeast and *Drosophila* cells ^28,30,32,42^, suggesting that the dependence on seipin for LD homeostasis is evolutionarily conserved.

**Figure 1.**
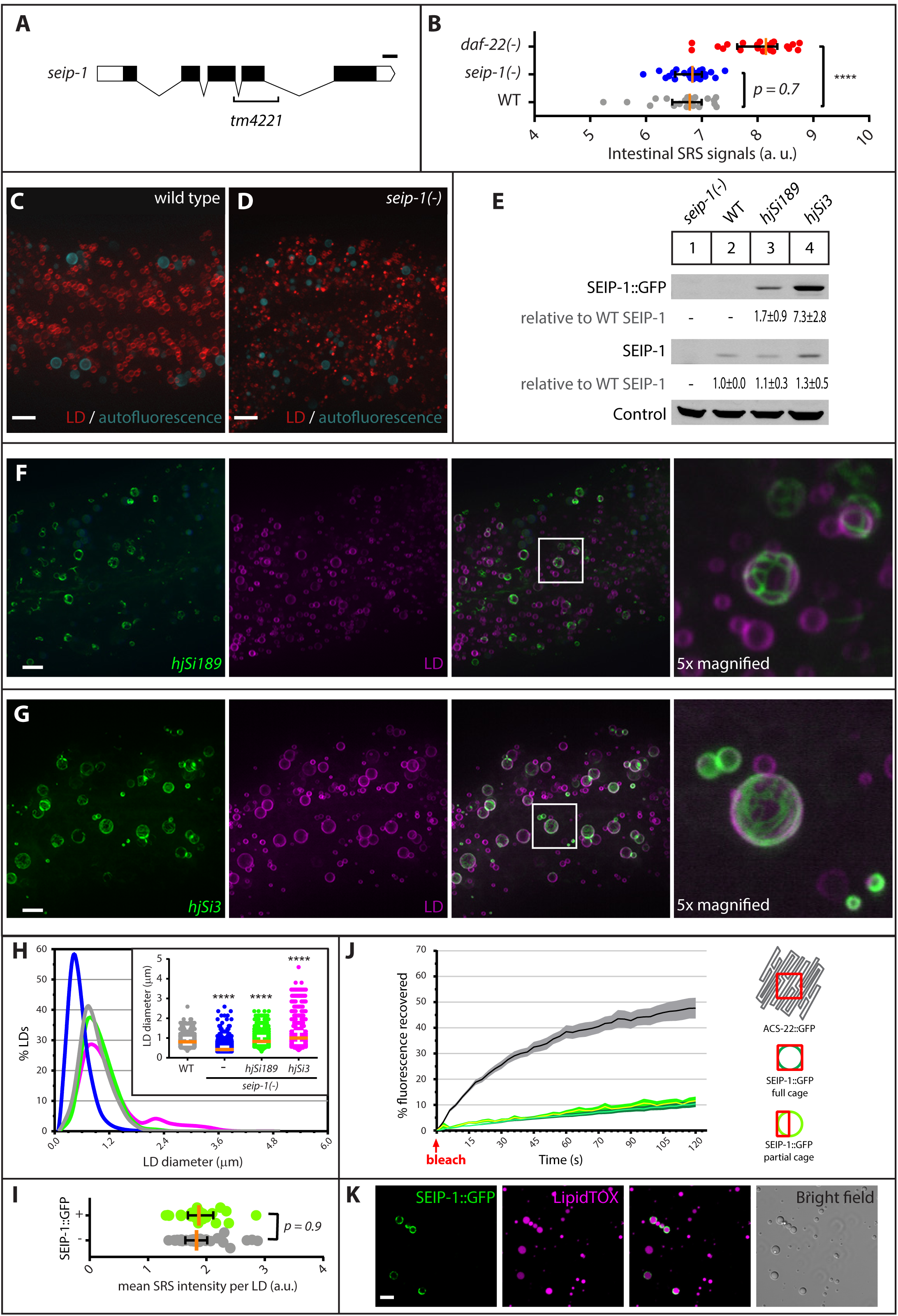
SEIP-1 regulates LD size from an ER subdomain. (A) Schematic representation of the *C. elegans* seipin ortholog, *seip-1*. Exons and untranslated regions are indicated by black and white boxes, respectively. The region deleted in the *tm4221* allele is indicated. Scale bar = 100bp. (B) Label-free quantification of neutral lipid content of individual animals by Stimulated Raman Scattering (SRS). A scatter plot summarizing the average intestinal SRS signals measured from at least 22 animals of each genotype is shown. Median with interquartile range is displayed. ****p < 0.0001 (unpaired t-test). (C) Visualization of LDs using the marker mRuby::DGAT-2 (*hjSi112*) in a wild-type (WT) larval L4 stage animal. mRuby is in red and autofluorescence from lysosome related organelles (LROs) is pseudocoloared cyan. A projection of 7.5μm z stack centering at the second intestinal segment is shown. (D) As in (C), but with a *seip-1 (tm4221)* mutant animal. (E) The expression levels of SEIP-1::GFP and endogenous SEIP-1 were determined by SDS-PAGE and immunoblotting with anti-SEIP-1 antibodies. A non-specific band at ∼95kDa served as a loading control. The expression level of endogenous SEIP-1 in wild type (WT) animals served as a reference for normalization of signals in other samples. The mean ± SD from at least three independent experiments is shown. (F) Visualization of SEIP-1::GFP (*hjSi189*, lane 3 in (E)) in a larval L4 stage animal. Autofluorescence is pseudocolored blue. The same LD marker was used as in (C) and the mRuby signal is pseudocolored magenta. A projection of 4.5μm z stack centering at the second intestinal segment is shown. The boxed area was magnified 5x and shown on the right. (G) As in (F), but with an animal over-expressing SEIP-1::GFP (*hjSi3*, lane 4 in (E)). (H) Frequency distribution of LD diameter. The curve was fitted twice using Fit Spline/LOWESS (20 points in smoothing window, 4000 segments) method based on a histogram with a bin size of 0.3μm (Figure S1D for an example). Inset: a scatter plot showing the median and inter-quartile range of LD size. Total number of LDs measured: WT = 1,680; *seip-1 (tm4221)* = 3,934; *hjSi189;seip-1 (tm4221)* = 2,224; *hjSi3;seip-1 (tm4221)* = 1,209. (I) Label-free quantification of lipid concentration of individual LDs by SRS. LDs with a diameter of 0.8-1.7μm in the second intestinal segment were selected for measurement. The scatter plot is formatted as in (B). Total number of LDs measured: SEIP-1::GFP (-) = 34; SEIP-1::GFP (+) = 22. (J) Percentage of recovered fluorescence of ACS-22::GFP (resident ER protein, *hjSi29*) and SEIP-1::GFP (*hjSi3*) in wild type animals. The connecting curves show the means and the filled areas show the SEM range. Number of photobleaching events: ACS-22::GFP = 17; SEIP-1::GFP full cage = 15; SEIP-1::GFP partial cage = 20. (H) Visualization of SEIP-1::GFP in LDs purified from *hjSi3* animals. LDs were stained with LipidTox Deep Red (pseudocolored magenta). For all fluorescence images, scale bar = 5μm.

To investigate how SEIP-1 affects LD size, we examined the sub-cellular localization of SEIP-1 by expressing a SEIP-1::GFP fusion protein at the endogenous level from a single copy transgene *hjSi189*, which is driven by the ubiquitous *dpy-30* promoter (Figure 1E, lane 3). The *hjSi189* transgene increased the median LD diameter of *seip-1(-)* animals to 0.83μm, which is similar to that in wild type animals (0.81μm) (Figure 1H, inset). We concluded that the SEIP-1::GFP fusion protein was functional. In the *C. elegans* intestine, we observed SEIP-1::GFP exclusively in nanotubes that formed rings and cages around a subset of LDs (Figure 1F). These nanotubes are most likely derived from ER tubules (see Figure 4). We named the SEIP-1::GFP positive structures peri-lipid droplet (peri-LD) cages. Similar observations were made when SEIP-1::tagRFP was expressed (Figure S1J). Furthermore, we used CRISPR to generate transgenic animals that express SEIP-1 with a 3xFLAG epitope tag at its C-terminus. Immunostaining revealed SEIP-1 positive signals around a subset of LDs (Figure S1H-I). Our results suggest that SEIP-1 exerts its effect at ER subdomains that are in close proximity to LDs.

### Over-expression of SEIP-1 increased LD size

The seipin orthologs have been found at ER-LD junctions ^27,30^. It has been proposed that the enrichment of seipin at these junctions promote the maturation of nascent LDs ^32^. Our observations in *C. elegans* suggested that SEIP-1 remained associated with a subset of mature LDs. To probe the functional significance of such association, we generated a single-copy transgene (*hjSi3*) that over-expressed a SEIP-1::GFP fusion protein specifically in the *C. elegans* intestine (Figure 1E, lane 4). The localization of SEIP-1::GFP to peri-LD cages persisted in these animals. Using mRuby::DGAT-2 to mark all LDs, we determined that approximately 40% of LDs were surrounded by peri-LD cages in each intestinal cell (Figure 1G and 2G).

**Figure 2.**
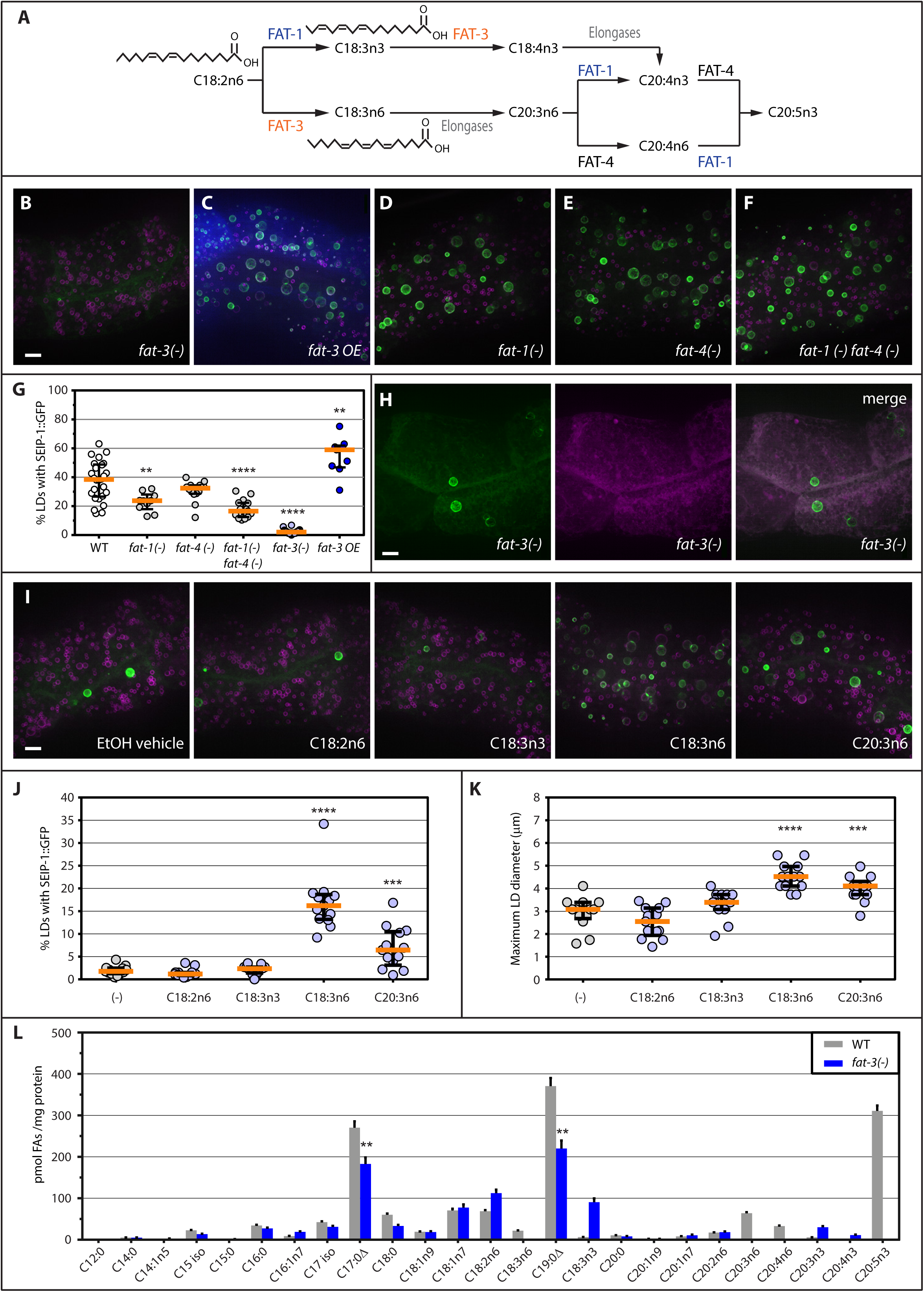
Mislocalization of SEIP-1::GFP in *fat-3* mutant animals. (A) The biosynthetic pathway for polyunsaturated fatty acids in *C. elegans*. (B) Visualization of SEIP-1::GFP (*hjSi3*) in a *fat-3 (ok1126)* mutant larval L4 stage animal. A projection of 4.5μm z stack centering at the second intestinal segment is shown. The LD marker mRuby::DGAT-2 (*hjSi112*) was used and the mRuby signal is pseudocolored magenta. (C) As in (B), but with a WT animal carrying a transgene that over-expressed FAT-3 (*Ex[fat-3p::fat-3 genomic DNA::SL2::tagBFP]*). (D-F) As in (B), but in *fat-1 (wa9), fat-4 (wa14)* or *fat-1 (wa9) fat-4 (wa14)* mutant backgrounds. (G) The percentage of LDs associated with SEIP-1::GFP. A scatter plot derived from at least 9 animals of each genotype is shown. Median with interquartile range is displayed. **p < 0.01; ****p < 0.0001 (unpaired t-test). (H) As in (B), but with a luminal ER mCherry marker (*hjSi158*) in a *fat-3 (ok1126)* mutant animal. The mCherry signal is pseudocolored magenta. (I) Visualization of SEIP-1::GFP (*hjSi3*) and mRuby::DGAT-2 (*hjSi112*) in a *fat-3(ok1126)* mutant larval L4 stage animals grown with either EtOH vehicle, linoleic acid (C18:2n6), α-linolenic acid (C18:3n3), γ-linolenic acid (C18:3n6) or dihomo-γ-linolenic acid (C20:3n6) supplementation. A projection of 4.5μm z stack centering at the second intestinal segment is shown. (J) The effects of PUFAs supplementation on the percentage of LDs associated with SEIP-1::GFP. A scatter plot derived from at least 11 animals of each condition is shown. Median with interquartile range is displayed. (K) As in (J), but the effects of PUFAs supplementation on the maximum LD diameter is shown. ***p < 0.001; ****p < 0.0001 (unpaired t-test). (L) The abundance of all fatty acids detected in WT or *fat-3 (ok1126)* larval L4 stage animals. Mean ± SEM of 4 independent samples is shown. **p < 0.01 (unpaired t-test). For all fluorescent images, scale bar = 5μm.

Opposite to the effect of loss of SEIP-1 activity on LD size, there was a significant increase in the median LD size in animals over-expressing SEIP-1::GFP (Figure 1H). This was in part due to the appearance of LDs >2μm in diameter, which were invariably associated with peri-LD cages. Nevertheless, SEIP-1 positive and negative LDs have similar concentration of neutral lipids, as determined by SRS (Figure 1I). Taken together, we conclude that over-expression of SEIP-1::GFP protein conferred a gain-of-function phenotype: expansion of a subset of LDs.

To further examine the spatial relationship between peri-LD cages with the ER tubular network, we generated animals that carry a second single-copy transgene that allowed the visualization of the ER lumen (signal sequence::mCherry::HDEL; *hjSi158*). Fluorescence signals from SEIP-1::GFP at the periphery of peri-LD cages juxtaposed with that from mCherry in the ER lumen (Figure S2K). However, the mobility of SEIP-1 was clearly different from a reference resident ER membrane protein, ACS-22/acyl-coA synthetase ^15^. Little recovery was observed when SEIP-1::GFP signals in whole or in part of cages were subjected to photo-bleaching (Figure 1J). This was in contrast to the rapid recovery of ACS-22::GFP in the general ER network. Next, we purified LDs from *hjSi3* transgenic animals by differential centrifugation and found that peri-LD cages labeled by SEIP-1::GFP remained stably associated with LDs (Figure 1K). Our imaging and biochemical results suggest that peri-LD cages are ER subdomains that preferentially associate with LDs. In addition, SEIP-1 is relatively immobile once incorporated into peri-LD cages, which may in part be due to regional differences of ER membrane constituents.

### Enrichment of SEIP-1 in peri-LD cages required polyunsaturated fatty acids

Next, we sought to determine the molecular mechanism for SEIP-1 enrichment in peri-LD cages. We mutagenized *hjSi3* animals with the chemical mutagen ethyl methane sulfonate (EMS) and obtained two mutants (*hj55* and *hj56*) in which SEIP-1::GFP was largely retained in the ER. Molecular cloning of these mutants identified mutations in the *fat-3* gene, which encodes a fatty acid desaturase for the synthesis of polyunsaturated fatty acids (PUFAs) in *C. elegans* (Figure 2A) ^43^. Specifically, FAT-3 introduces a double bond to C18:2n6 and C18:3n3 to yield C18:3n6 and C18:4n3, respectively. We noted that aberrant localization of SEIP-1::GFP was also observed in mutants that harbored a deletion in the *fat-3* gene (*ok1126*) (Figure 2B). This was not due to a change in the expression level of endogenous SEIP-1 and SEIP-1::GFP (Figure S2A, lane 7), nor a change in LD size distribution of *fat-3(-)* animals (Figure S2B). We concluded that proper SEIP-1::GFP targeting is dependent on PUFAs that are direct or indirect products of the FAT-3 desaturase. Since FAT-3 does not co-localize with SEIP-1 and is distributed throughout the ER network (Figure S2C-D), it is plausible that additional mechanisms may be used to concentrate FAT-3 products in ER subdomains.

Based on previous reports, FAT-3 is required for the biosynthesis of multiple PUFAs ^43^. This was confirmed using a quantitative gas chromatography-mass spectrometry (GCMS) based assay (Figure 2L). Does SEIP-1::GFP enrichment in peri-LD cages require specific species of PUFAs? To address this question, we examined the localization of SEIP-1::GFP in *fat-1*, *fat-4* and *fat-1 fat-4* loss-of-function mutant animals. FAT-1 and FAT-4 encode fatty acid desaturases that act upstream and downstream of FAT-3 (Figure 2A). The percentage of LDs associated with SEIP-1::GFP was significantly reduced in *fat-1* and *fat-1 fat-4* mutant animals although the effect was not as pronounced as in *fat-3* animals (Figure 2D-G). Again, no change in the expression level of endogenous SEIP-1 and SEIP-1::GFP was observed (Figure S2A, lanes 5 and 6). When FAT-3 was over-expressed, the percentage of LDs associated with SEIP-1::GFP was increased, suggesting that FAT-3 products play an instructive role in SEIP-1 targeting (Figure 2C, 2G). Is a specific FAT-3 product required? To this end, we supplemented FAT-3 substrates (C18:2n6 and C18:3n3) or products (C18:3n6 and C20:3n6) to the *E. coli* diet of *fat-3(-)* animals. The enrichment of SEIP-1::GFP in peri-LD cages was significantly increased by C18:3n6 and C20:3n6 supplementation, while C18:2n6 or C18:3n3 had no effect (Figure 2I and J). This was accompanied by an increase in the maximal LD diameter in C18:3n6 and C20:3n6 treated animals (Figure 2K), consistent with the role of SEIP-1 in promoting LD expansion. Taken together, our genetic and dietary supplementation experiments indicate that proper targeting of SEIP-1::GFP is at least partially dependent on specific PUFAs.

### Enrichment of SEIP-1 in peri-LD cages required microbial fatty acids

Our quantitative GC-MS analysis revealed that in addition to a reduction of PUFAs, the abundance of two cyclopropane fatty acids (CFAs), C17:0Δ and C19:0Δ, was also significantly reduced in *fat-3(-)* animals (Figure 2L). Since CFAs are strictly derived from *E. coli* and are not synthesized by *C. elegans*, our results implied that *fat-3(-)* animals are partially defective in absorbing dietary microbial fatty acids. CFAs are synthesized by the CFA synthase from monounsaturated fatty acids (Figure S3A) ^44^. They are major constituents of bacterial phospholipids that are important for membrane integrity under stress ^45^. We next asked if the mis-localization of SEIP-1::GFP in *fat-3(-)* animals was in part due to a reduction of diet-derived CFAs. To this end, we obtained the *E. coli* CFA synthase deletion mutant and its parental strain BW25113 from the Keio collection ^46^. Using GC-MS, we confirmed that the *Δcfa* strain indeed was unable to synthesize C17:0Δ and C19:0Δ (Figure S3B). Next, we fed *hjSi3; hjSi112* transgenic animals that expressed SEIP-1::GFP and mRuby::DGAT-2 with BW25113 and *Δcfa E. coli*. Using mRuby::DGAT-2 as a pan-LD marker in the intestine, we noted a reduction of median LD diameter and total LD volume in animals fed *Δcfa E. coli* (Figure 3C, S3C). Furthermore, the percentage of LDs that are associated with SEIP-1::GFP peri-LD cages was also significantly reduced (Figure 3D). Quantitative GC-MS analysis indicated an almost complete loss of C17:0Δ and C19:0Δ, and a concomitant increase of C16:1n7 and C18:1n7 in the fatty acid profile of worms fed *Δcfa E. coli* (Figure 3E). This was consistent with C16:1 and C18:1n7 being the precursors of C17:0Δ and C19:0Δ in *E. coli*, respectively (Figure S3B). Our results demonstrate that *C. elegans* obtain CFAs from their microbial diet and incorporate them efficiently into their cellular lipids. Such dietary CFAs have a significant impact on fat content, LD morphology and SEIP-1 targeting.

**Figure 3.**
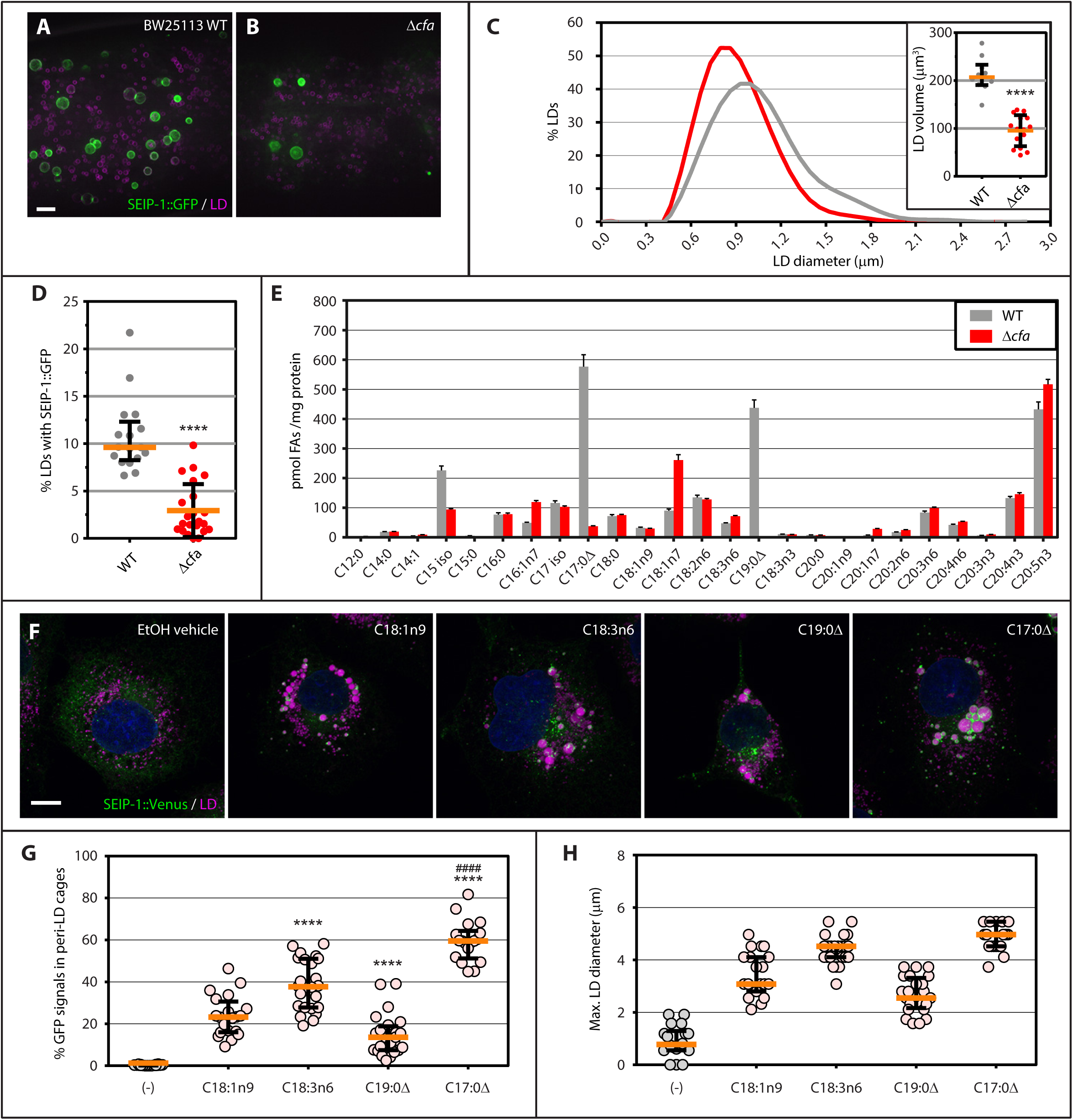
Bacterial cyclopropane-FAs regulate SEIP-1::GFP targeting. (A) Visualization of SEIP-1::GFP (*hjSi3*) in a larval L4 stage animal fed on wild type (WT) *E. coli BW25113*. A projection of 4.5μm z stack centering at the second intestinal segment is shown. The LD marker mRuby::DGAT-2 (*hjSi112*) was used and the mRuby signal is pseudocolored magenta. Scale bar = 5μm. (B) As in (A), but with an animal that was fed *E. coli BW25113 Δcfa*. (C) Frequency distribution of LD diameter in larval L4 stage animals that were fed wild type (grey) or *Δcfa* (red) *E. coli*. The curve was fitted twice using Fit Spline/LOWESS (20 points in smoothing window, 4000 segments) method based on a histogram with a bin size of 0.3μm. Total number of LDs measured: *BW25113* WT fed = 2,414; *BW25113 Δcfa* fed = 2,704. A scatter plot summarizing the total LD volume in the second intestinal segment of each animal is shown in the inset (n ≥ 10). Median with interquartile range is displayed. ****p < 0.0001 (unpaired t-test). (D) The lack of dietary CFA reduced the percentage of LDs associated with SEIP-1::GFP. A scatter plot derived from at least 17 animals on each diet is shown. Median with interquartile range is displayed. ****p < 0.0001 (unpaired t-test). (E) The abundance of all fatty acids detected in worms that were fed wild type (grey) or *Δcfa* (red) *E. coli*. Mean ± SEM of 4 independent samples is shown. (F) COS7 cells stably expressing SEIP-1::Venus were supplemented with either EtOH vehicle, oleic acid (C18:1n9), γ-linolenic acid (C18:3n6), phytomonic acid (C19:0Δ) or cis-9,10-methylenehexadecanoic acid (C17:0Δ). Blinded experiments were performed: the identity of FAs supplemented in each dish was not known to the experimenter who took the images. LDs were stained with LipidTox Red (pseudocolored magenta) and nuclei were stained with Hoechst 33342 (blue). A projection of 4.5μm z stack is shown. Scale bar = 10μm. (G) The effect of FA supplementation on the percentage of SEIP-1::Venus signals in peri-LD cages. A scatter plot derived from at least 18 cells of each condition is shown. Median with interquartile range is displayed. (H) As in (G), but the effect of FA supplementation on the maximum LD diameter is shown. ****p < 0.0001 (unpaired t-test against C18:1n9); ^####^p<0.0001 (unpaired t-test against C18:3n6).

### PUFAs and CFAs regulate SEIP-1 targeting in mammalian cells

To investigate if SEIP-1 localizes to peri-LD structures in mammalian cells, we generated a COS7 cell line that stably expressed a SEIP-1::Venus green fluorescent fusion protein. When these cells were loaded with oleic acid (C18:1n9), SEIP-1::Venus was enriched in peri-LD structures, similar to what we observed in *C. elegans* (Figure 3F). Using the same cells, we next tested the effect of PUFAs and CFAs on SEIP-1::Venus targeting. In comparison with oleic acid (C18:1n9), supplementation of C18:3n6 and C17:0Δ had significantly stronger effects on promoting LD expansion, which correlated with SEIP-1::Venus enrichment in peri-LD cages (Figure 3F-H). Furthermore, the effect of C17:0Δ was dose-dependent (Figure S3D-F). Our results suggest that polyunsaturated C18:3n6 and microbial C17:0Δ play conserved roles in regulating SEIP-1 targeting in mammalian cells.

### Conservation of seipin function and localization in C. elegans and mammalian cells

In COS7 cells that stably expressed SEIP-1::Venus and treated with oleic acid, LDs marked by SEIP-1::Venus were significantly larger than those that were not marked in the same cells (Figure 4A-B). Similar observations were made in HeLa cells (Y.H., unpublished data). Therefore, the ability of SEIP-1 to increase the size of its associated LDs appeared to be conserved in *C. elegans* and mammalian cells. Next, we expressed the human seipin protein fused to GFP from a single-copy transgene in *C. elegans*. The human seipin::GFP fusion protein was functional since it restored the median LD size of *seip-1(-)* animals to wild type level (Figure 4C). Similar to *C. elegans* SEIP-1, human seipin was found to associate with a subset of LDs in tubular structures (Figure 4D). Taken together, the mechanism by which seipin family members regulate LDs from peri-LD cages is conserved.

**Figure 4.**
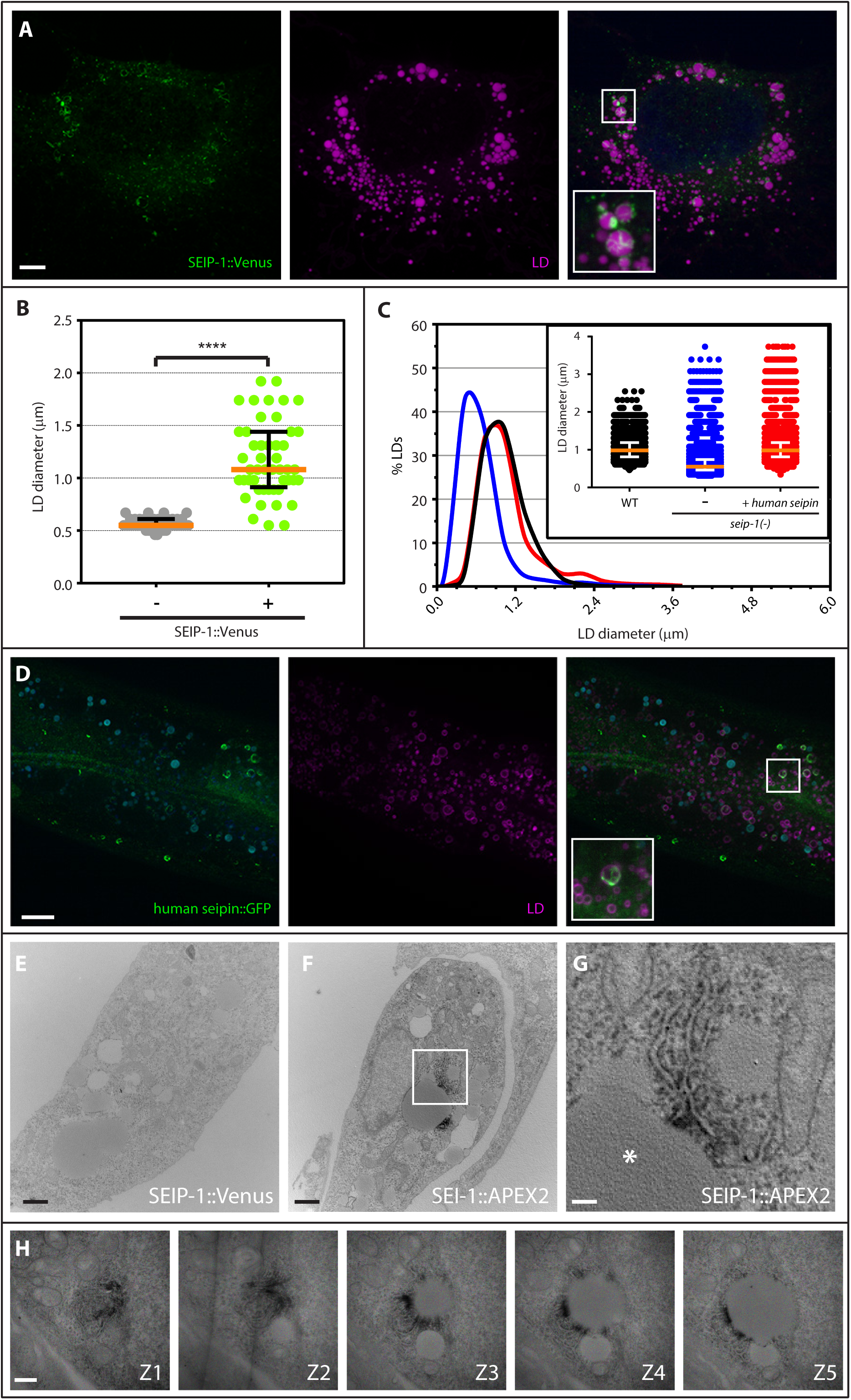
Enrichment in peri-LD cages is an evolutionarily conserved property of seipin. (A) The SEIP-1::Venus protein was stably expressed in COS7 cells. The cells were incubated with oleic acid prior to fixation. A projection of 4.5μm Z stack is shown. The boxed area was magnified 3x and shown in the inset. Scale bar = 5μm. (B) Quantification of LD size in COS7 cells stably expressing SEIP-1::Venus. A scatter plot of data from 48 LDs of each category is shown. Median with interquartile range is displayed. ****p < 0.0001 (unpaired t-test) (C) Frequency distribution of LD diameter in larval L4 stage animals of indicated genotypes. The human seipin isoform 2 was expressed from a single-copy transgene (*hjSi223*). The curve was fitted twice using Fit Spline/LOWESS (20 points in smoothing window, 4000 segments) method based on a histogram with a bin size of 0.3μm. Inset: a scatter plot showing the median and inter-quartile range of LD size. Total number of LDs measured: WT = 4,600; *seip-1(tm4221)* = 6,069; *seip-1(tm4221);hjSi223* = 4,258. (D) Visualization of human seipin::GFP in a *seip-1 (tm4221)* mutant larval L4 stage animal. A projection of 3μm z stack centering at the second intestinal segment is shown. The LD marker mRuby::DGAT-2 (*hjSi112*) was used and the mRuby signal is pseudocolored magenta. Autofluorescence is pseudocolored blue. The boxed area was magnified 2.5x and shown in the inset. Scale bar = 10μm. (E) Transmission electron micrograph of a COS7 cell stably expressing SEIP-1::Venus protein. The cell was incubated with oleic acid prior to fixation, and stained with diaminobenzidine (DAB), uranyl acetate, and osmium tetroxide prior to electron microscopy. No dark deposits were detected in this negative control. Scale bar = 1μm. (F) As in (E), but with SEIP-1::V5-APEX2 fusion protein stably expressed. (G) The boxed area in (F) was magnified 5x. Dark deposits indicate the localization of SEIP-1::V5-APEX2 at ER tubules. The LD is indicated by an asterisk. Scale bar = 0.2μm. (H) As in (G), but with five consecutive sections (150nm thick) shown. Scale bar = 0.5μm

### Peri-LD cages are ER subdomains

We postulated that peri-LD cages are composed of ER tubules, based on their morphology and their close proximity and tight association with LDs. To verify our hypothesis, we generated COS7 cells that stably expressed a SEIP-1::APEX2 fusion protein at low levels, using the Sleeping Beauty Transposon system ^47,48^. The SEIP-1::APEX2 fusion protein could be detected around enlarged LDs, suggesting the function of SEIP-1 was preserved (Figure S4A). The APEX2 peroxidase reacts with diaminobenzidine (DAB) to generate local deposits that can be stained with uranyl acetate and osmium tetroxide in electron microscopy. Accordingly, we found that SEIP-1::APEX2 was highly enriched on ER tubules in the proximity of LDs in oleic acid loaded cells (Figure 4F-H, S4D-K). The APEX2 mediated dark deposits were not detected in negative control cells that expressed SEIP-1::Venus (Figure 4E). The SEIP-1::APEX2 positive ER tubules showed similar morphology to those that are enriched with enzymes in the TAG biosynthetic pathways in *Drosophila* cells ^12^. It has been proposed that such ER tubules constitute direct connections between ER and LDs. Our results suggest that peri-LD cages are ER subdomains that enable the assembly of proteins in support of LD expansion.

## Discussion

In this paper, we combined genetic, biochemical, light and electron microscopy approaches to demonstrate the preferential enrichment of SEIP-1/seipin to an ER subdomain, termed the peri-LD cage. Peri-LD cages selectively envelope a subset of LDs that have high propensity for expansion. Recruitment of SEIP-1 to peri-LD cages demands PUFAs and microbial CFAs. More specifically, γ-linolenic acid (C18:3n6) and cyclopropane fatty acid (C17:0Δ) have the strongest effect on SEIP-1 targeting in *C. elegans* and mammalian cells. Our results suggest that the heterogeneity of LDs can be enforced not only by LD proteins, but through inter-organelle contacts with the ER.

How does the enrichment of SEIP-1 in peri-LD cages increase LD size? In the energy-replete state, a balance of lipolysis and TAG synthesis determines LD size ^12,49^. Attenuation of lipolysis, i.e. blocking the release of fatty acids from triglycerides in LDs, will therefore increase LD size. Lipolysis is rate-limited by the recruitment of adipose triglyceride lipase (ATGL) to the LD surface ^50^. Accordingly, large LDs accumulate in non-adipose tissues in ATGL deficient humans ^51,52^. We have previously shown that the *C. elegans* ATGL ortholog (ATGL-1) is found at LDs ^40^. Since the ATGL-1::GFP fusion protein was observed at the surface of LDs with or without peri-LD cages (Figure S4L-N), we concluded that peri-LD cages did not impede lipolysis by blocking ATGL-1 recruitment. Development of assays for measuring the lipolytic rate at the single LD level will be necessary to probe additional effects of peri-LD cages on lipolysis.

The de novo synthesis of triglycerides (TAG) is dependent on LD associated DGAT2. In mammalian cells, DGAT2 transits from the ER to the LD surface upon lipid loading ^12,18^. In *C. elegans*, the DGAT2 ortholog (DGAT-2) associates stably with LDs ^15^. Throughout this study, mRuby::DGAT-2 was used to mark all LDs and we did not observe enhanced DGAT-2 residency at LDs with peri-LD cages. Nevertheless, it is plausible that peri-LD cages can increase the local concentration of additional proteins that are rate-limiting for TAG synthesis. These proteins should be well conserved since peri-LD cage association increased LD size in both *C. elegans* and mammalian cells. For example, the Glycerol-3-phosphate acyltransferase 4 (GPAT4), which acts upstream of DGAT2 in the TAG synthesis pathway, is also known to migrate from the ER to LDs ^12^. Although the mammalian GPAT4 shares sequence homology with a number of *C. elegans* acyltransferases, its functional ortholog awaits to be identified definitively.

In budding yeast, several proteins including Ldb16 and Ldo45 have been shown to act with the seipin ortholog Fld1 at ER-LD junctions to regulate LD size ^53–58^. However, BLAST homology searches for their orthologs in *C. elegans* did not return significant hits. Since the conserved central loop region of seipin orthologs is flanked by divergent N- and C-termini, it is tempting to speculate that seipin orthologs may have conserved and species-specific partners that modulate LD morphology in a convergent manner.

Cellular membranes are in large part composed of phospholipids that consist of a head group and two fatty acyl chains. The combination of saturated, monounsaturated, and polyunsaturated acyl chains in phospholipids modulates the packing and curvature of membranes that contribute to organelle identity and function ^34^. The well-established importance of polyunsaturated fatty acids (PUFAs) for neurons has recently been linked to their enrichment in membranes of synaptic vesicles and photoreceptor discs ^59–61^. Although humans obtain PUFAs from their diet, PUFAs are synthesized endogenously by a set of desaturases in *C. elegans* ^43^. Based on biochemical and molecular dynamics simulations, PUFAs have the unique ability to support the formation of highly curved and densely packed membranes ^62^. It is plausible that ER tubules that constitute peri-LD cages are composed of such membranes since SEIP-1 was immobile once incorporated into peri-LD cages (Figure 1J). This could be due to the tight fitting of fatty acyl chains around SEIP-1. PUFAs in peri-LD cages may also act as diffusion barriers to additional proteins, such as those that support LD expansion. Our work supports the idea that PUFAs have fundamental structural roles in defining membrane territories.

There is an increasing appreciation on how the host metabolism can be modulated by the microbiome, in part through microbial metabolites ^63,64^. Our work reveals cyclopropane fatty acids (CFAs, C17:0Δ and C19:0Δ) as new players in host-microbe interaction. Cyclopropane fatty acyl chains are generated through the action of CFA synthase on monounsaturated precursors in membrane phospholipids of *E. coli* and other bacteria ^44^. Since there is no eukaryotic CFA synthase, the detection of CFAs in *C. elegans* indicates their effective absorption and incorporation into host cellular lipids. CFAs have also been detected in human adipose tissue where seipin is strongly expressed ^22,36^. Nevertheless, it was hitherto unclear how CFAs play a role in host metabolism. Our work hinted at one possible mechanism: CFA-dependent protein targeting. We noted that worms fed CFA deficient *E. coli* have less fat and smaller LDs (Figure S3C). These animals also grow slower than those that are fed a CFA replete diet (C.Z., unpublished data). Therefore, microbial CFAs may have broad effects on host physiology, similar to PUFAs.

In summary, our results reveal unexpected and conserved roles of microbial and polyunsaturated fatty acids on cellular fat storage. They modulate SEIP-1/seipin targeting to peri-LD cages and in turn promote LD expansion in *C. elegans* and mammalian cells. The selective association of peri-LD cages to a subset of LDs also strengthen the notion of LD heterogeneity ^10^. Distinct populations of LDs may carry specific cargos beyond neutral lipids. They may be subject to differential turnover during fed or fasted states. Our results implicate the selective deployment of dietary and endogenous fatty acids in defining organelle identity through specialized membrane environments.

## Methods

### Strains and Transgenes

The wild type *C. elegans* strain was Bristol N2. All experimental animals were maintained at 20°C. The following alleles were obtained from the *Caenorhabditis* Genetics Center (CGC), which is funded by NIH Office of Research Infrastructure Programs (P40 OD010440): LG III, *unc-119 (ed3)*; LG X, *cept-1 (et10), cept-1 (et11)*; LG IV, *fat-1 fat-4 (wa9 wa14), fat-3 (ok1126)*. The *seip-1(tm4221)* allele was obtained from Dr. Shohei Mitani (National Bioresource Project for the nematode). The following transgenes were used:

hjSi3[vha-6p::seip-1 cDNA::GFP_TEV_3xFLAG::let-858 3’ UTR] II
hjSi30[vha-6p::seip-1 cDNA(A185P)::GFP_TEV_3xFLAG::let-858 3’ UTR] II
hjSi112[vha-6p::3xFLAG_TEV_mRuby2::dgat-2 cDNA::let-858 3’UTR] IV
hjSi189[dpy-30p::seip-1 cDNA::GFP_TEV_3xFLAG::tbb-2 3’ UTR] II
hjSi222[fat-3p::fat-3 genomic DNA::GFP-TEV-3xFLAG::fat-3 3’ UTR] I
hjSi434[dpy-30p::seip-1 cDNA::tagRFP::tbb-2 3’ UTR] II
hjEx23[fat-3p::fat-3 genomic DNA_SL2_tagBFP2::fat-3 3’UTR]
dhs-3(hj120) [dhs-3::GFP_3xFLAG]
atgl-1(hj91) [atgl-1::GFP_3xFLAG]
seip-1(hj196) [seip-1::TEV_3xFLAG]

### C. elegans Genetic Screen

To isolate mutant animals with altered localization of SEIP-1::GFP, we mutagenized *hjSi3* animals with ethyl methane sulfonate (EMS) using standard procedures. We screened ∼38,000 haploid genomes and selected mutant F2 animals on a UV fluorescence dissecting microscope (Leica). Mutant animals were back-crossed with *hjSi3* animals at least twice and their phenotype was subsequently verified by confocal microscopy. For molecular cloning, we focused on one complementation group comprised of *hj55* and *hj56* alleles. Genetic mapping with the Hawaiian isolate CB4856 placed both alleles on LGIV. Molecular lesions in the *fat-3* gene were identified by whole-genome sequencing (Illumina) and verified by targeted Sanger sequencing. The *hj55* and *hj56* alleles encode the mutations Gln325Stop and Gly329Asp, respectively. The *fat-3(ok1126)* deletion allele, outcrossed four times with N2, was subsequently used as the reference strain. The mis-localization of SEIP-1::GFP in the *hjSi3; fat-3(ok1126)* strain could be rescued by a *fat-3p::fat-3 genomic DNA::SL2::tagRFP::fat-3 3’ UTR* transgene.

### Fluorescence Imaging of C. elegans and mammalian cells

Fluorescence imaging was performed as described ^19^. The following changes in acquisition parameters were made. Fluorescence images of L4 larval animals or mammalian cells were acquired on a spinning disk confocal microscope (AxioObeserver Z1, Carl Zeiss) equipped with a piezo Z stage using a 100x, numerical aperture (NA) 1.46 oil Alpha-Plan-Apochromat objective, on an Neo sCMOS camera (Andor) controlled by the iQ3 software (Andor). For GFP, a 488nm laser was used for excitation and signals were collected with a 500-550nm emission filter. For mCherry and tagRFP, a 561nm laser was used for excitation and signals were collected with a 580.5-653.5nm emission filter. For autofluorescence from lysosome-related organelles, a 488nm laser was used for excitation and signals were collected with a 580.5-653.5nm emission filter. For tagBFP, a 405nm laser was used for excitation and signals were collected with a 417-477nm emission filter. Optical sections were taken at 0.5μm intervals and z stacks of 8μm-10μm were exported from iQ3 to Imaris 8 (Bitplane) for processing. For each image, 10 planes in total (4.5μm in z axis) were used for 3D-reconstruction and measurement of LD diameter unless otherwise indicated.

Photo-bleaching experiments were performed on larval L4 stage animals using the spinning disk confocal microscope described above. The region of interest (ROI) was chosen to cover the entire or half peri-LD cage in *hjSi3* animals. For each FRAP event, the size of the ROI was kept constant for both background and reference regions. One or more ROIs of at least eight animals were examined for each strain. Normalized fluorescence intensity (NI) was calculated using the following formula, taking background fluorescence (Back) and fluorescence in an unbleached reference ROI (Ref) into consideration.

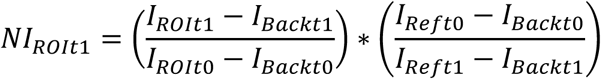

### Label-free Lipid Imaging in C. elegans

For analyzing single lipid droplets, an integrated femtosecond (fs) stimulated Raman scattering (SRS) and two-photon excited fluorescence (TPEF) microscope system was used for label-free imaging of LDs in *C. elegans* ^39^. L4 larval animals (*hjSi3*) were dissected in the physiological buffer (150mM NaCl, 3mM KCl, 2mM MgCl_2_, 3mM CaCl_2_, 10mM glucose and 15mM HEPES; 340-345mmol/kg) containing 0.2mM levamisole and mounted on a 2% agarose pad. Raw signals from GFP, SRS and tryptophan were acquired sequentially from a field of view (FOV) of 30 × 30 μm in z stacks of 7μm at 0.5μm intervals. For quantification, an averaged background noise 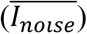 from photodiode and lock-in amplifier (LIA) was measured and SRS signals of glycerol (*I_glycerol_*) in the same FOV were acquired for normalization prior to acquisition of the GFP, SRS and tryptophan signals. LD in dissected *hjSi3* animals that was 0.8-1.7μm in diameter and present in focal plane 1-8 (3.5μm in z axis) was used for quantification. For each LD in each focal plane, total intensity of normalized SRS signals (*I_plane_*) was calculated by:

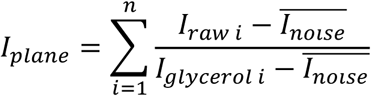

where *I_raw i_* is the intensity of *i^th^* pixel present in the LD and *n i*s the number of all pixels enclosed in the LD. *I_plane_* from all focal planes that each LD spanned were summed up as the total intensity of each LD (*TI_LDs_*). Volume of each LD was calculated using ImageJ in raw SRS images.

For whole-worm lipid measurement, live L4 larval animals were directly immersed in the physiological buffer containing 0.2mM levamisole and mounted on a 8% agarose pad. For each animal, a projection image with maximal SRS intensity was acquired using a picosecond SRS microscope system as previously developed ^65^ equipped with a 20x air objective (Plan-Apochromat, 0.8 NA, Zeiss). Quantification was performed according to published protocols ^66^. Briefly, raw total intestinal SRS intensity of each animal was adjusted by 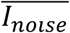 and *I_glycerol_* as described above, and subsequently averaged by the total intestinal pixel number. At least 20 animals were imaged and quantified for each genotype.

### Antibodies and Western Blotting for C. elegans

Antibodies against SEIP-1 were raised by injection of the peptide ^261^KKEEPGLLDLRKRK, corresponding to the C-terminus of SEIP-1, into rabbits (YenZym). Antibodies were purified against the antigen immobilized on the AminoLink Plus Coupling resin (Pierce, #44894) and used in Western blotting at a dilution of 1:100. IRDye 800CW donkey anti-rabbit secondary antibodies were used at 1:5000. Fluorescence signals were visualized using an Odyssey Infrared Imaging system (LI-COR) and analyzed using Odyssey software according to manufacturer’s instructions. Background signals were measured from multiple areas of the blot without proteins, averaged, and subtracted from integrated intensity of each protein band. The non-specific signal from a ∼95kDa protein was used as the loading control unless otherwise indicated.

### LDs Isolation from C. elegans

LDs isolation from *C. elegans* was performed using a published method ^67^. For fluorescence imaging, 10μl LD sample was diluted with 90μl buffer B (20mM HEPES, 100mM KCl, 2mM MgCl2, pH 7.4) and incubated with 1:1000 diluted LipidTox Deep Red (ThermoFisher, #H34477) for 30min on ice. Fluorescence images were acquired using a confocal microscope (Olympus, FV1000).

### Dietary Supplementation of Polyunsaturated Fatty Acids (PUFAs)

Pure PUFAs: linoleic acid (LA), α-linolenic acid (ALA), γ-linolenic acid (GLA) and dihomo-γ-linolenic acid (DGLA) (Nu-Chek Prep) were dissolved in absolute ethanol at a final concentration of 50mM as stock solutions. Each stock solution was freshly diluted in sterilized Milli-Q water (30μl 50mM PUFAs and 120μl water) and overlaid on top of a seeded bacteria lawn in a 6cm nematode growth media (NGM) plate. The plates were immediately dried in a dark laminar flow hood for 30min. For fluorescence imaging, six *hjSi3;hjSi112* or *hjSi3;hjSi112 ok1126* L4 larvae were transferred onto each PUFAs-supplemented plate and allowed to lay eggs for 2 days. Two replicates were performed for each strain in each PUFAs-supplemented condition. For each replicate, 6-8 of their progenies at the L4 larval stage were imaged.

### Generation of stable SEIP-1::Venus and SEIP-1::V5-APEX2 cell lines

COS7 cells stably expressing SEIP-1::Venus or SEIP-1::V5-APEX2 were generated by the Sleeping Beauty (SB) transposon system ^47,48^. Cells were co-transfected with pCMV(CAT)T7-SB100 that bore the SB Transposase and a plasmid based on pSBbi-Hyg (SEIP-1::Venus) or pSBbi-RH (SEIP-1::V5-APEX2) using Lipofectamine 2000 (Life Technologies). Selection of stable integrants was facilitated by 0.5mg/mL hygromycin for at least 7 days. The resulting cell population was further sorted using a Becton Dickinson Influx system and arbitrarily sub-divided into “Low”, “Medium” and “High” fractions based on the fluorescence intensity of Venus (for SEIP-1::Venus cells) or TdTomato (for SEIP-1::V5-APEX2 cells).

### Transmission electron microscopy

Chemical staining and dehydration of COS7 cells that stably expressed SEIP-1::V5-APEX2 were performed based on published methods ^48^. Prior to fixation, cells were incubated with 400μM oleic acid-BSA complex (Sigma) for 24h. Fixation was conducted using 2.5% glutaraldehyde (Electron Microscopy Science) in wash buffer (100mM sodium cacodylate with 2mM CaCl2, pH 7.4). For diaminobenzidine (DAB) staining, 5mg/mL DAB free base (Sigma) stock solution dissolved in 0.1M HCl was freshly diluted with 10mM H_2_O_2_ in cold wash buffer, and cells were incubated in the solution for 10min. SEIP-1::Venus stable cells were used as the negative control for assessing non-specific DAB deposition. 2% osmium tetroxide (Electron Microscopy Science) staining was performed for 40min in cold wash buffer, followed by incubation with chilled 2% uranyl acetate in ddH2O overnight at 4°C. Infiltration was performed at room temperature with a progressive series (10%, 25%, 50%, 75% and 100%) of EMBED-812 resin to ethanol dilutions. Specimens were incubated for at least 3 hours in each dilution. After three changes of 100% Epon resin, samples were then transferred to a plastic mold and cured at 60°C for 40h ^68^. Cured blocks were then trimmed and sectioned with a diamond knife. 150nm thin sections were cut and mounted on copper slot grids coated with formvar and further imaged with a Hitachi H-7650 transmission electron microscope operated at 80KV.

For 3D model reconstruction, serial images were stacked and aligned using IMOD software package (Boulder Laboratory of 3D Electron Microscopy of the Cell, University of Colorado at Boulder). 3D modeling of ER without obvious DAB/APEX2 staining and LDs were performed using 3dmod graphic module while DAB/APEX2 precipitates were reconstructed using the auto-contour function under high contrast with a threshold level.

### Immunostaining endogenous SEIP-1 in dissected C. elegans

Animals were dissected, fixed and immunostained as according to published methods with minor modifications. Briefly, ∼200 L4 larval animals of each genotype were dissected in 1xPBS/0.0005% Triton X-100 (Sigma) with 0.2mM levamisole for intestine extrusion and fixed for 4h at 4°C in 2% paraformaldehyde. At room temperature, the fixed animals were permeabilized for 1h in 1xPBS/0.1% Triton X-100, followed by a 1h blocking in 1xPBS/0.1% Tween20/1% bovine serum albumin (BSA). Primary (1:100 diluted monoclonal anti-FLAG antibody (M2) (Sigma)) and secondary antibody (1:100 diluted Alexa Fluor 488 Donkey anti-Mouse IgG (H+L) (Invitrogen)) incubations were both performed overnight at 4°C in 1xPBS/0.1% Tween20/1% BSA. Sample was post-stained by HCS LipidTOX™ red neutral lipid stain (Invitrogen) and Hoechst 33342 (Invitrogen) according to manufacturer’s instructions and mounted onto a 35mm MatTek glass bottom dish coated with poly-D-lysine prior to imaging.

### Total lipid extraction and FAMEs preparation for gas chromatography-mass spectrometry (GC-MS) analysis

Sample was prepared and extracted according to published methods ^69^ with slight modifications. For total lipid extraction from bacteria, ∼15mL overnight culture was concentrated 10x and spread on a 10cm NGM plate. The plate was immediately dried in a laminar flow hood and incubated at 20°C for 2 days. Bacteria was rinsed off the plate, pelleted and washed 3 times using 0.9% NaCl solution. Two plates were combined as one sample, at least three samples were analyzed for each bacterial strain. After the last wash, the pellet was re-suspended and sonicated in ice water bath for 5 times at 40 % amplitude with a 2min on/30sec off cycle. 1/10 of the lysates were sampled to determine the soluble protein concentration using the BCA Protein Assay Kit (Pierce) and used for subsequent data normalization. Known amount of palmitic acid-D31 (Sigma) was added to the remaining lysates in order to control for extraction efficiency, sample loss and subsequent transmethylation efficiency. One volume of the remaining lysates were subsequently extracted using ten volumes of pre-chilled Folch solution (chloroform : methanol, 2:1, v:v) followed by five volumes of chloroform. Total lipids were pooled together from bottom chloroform phases into a clean glass vial with a PTFE cap and dried under a gentle stream of nitrogen at 37°C.

For FAs analysis of *C. elegans*, ∼4000 newly hatched L1 larvae were seeded onto each plate prepared as described above and allowed to grow to the late-L4 stage. L4 larvae were harvested and washed four times with 1xPBS/0.001% Tween 20. Worm pellets were snap-frozen and stored in −80°C prior to lipid extraction. Each plate of worms was used as one sample, at least 4 independent samples were used for each condition. For lipid extraction, 500μl of 1xPBS/0.001% Tween 20/ 0.001% butylated hydroxytoluene (BHT) was added to each sample pellet and sonicated in ice water bath for 15 times at 40 % amplitude with a 2min on/30sec off cycle. 100μl of the sonicated sample was mixed with an equal volume of 4 x SDS buffer (8% SDS, 200mM Tris-HCl pH 6.8) and denatured at 95°C for 10min. Concentration of soluble protein was determined using the BCA Protein Assay Kit (Pierce) and used for subsequent data normalization. Known amount of heptadecanoic acid (C17:0, Nu-Chek Prep) was added to the remaining lysates in order to control for extraction efficiency, sample loss and subsequent trans-methylation efficiency. The remaining 400μl worm lysate was subsequently subjected to total lipid extraction as described for bacteria above.

Dried lipids were re-dissolved in 1.95mL methanol. Trans-methylation was carried out by adding 50μL sulfuric acid and incubating the sample mixture at 80°C for 1h. After adding 1.5mL Milli-Q water, fatty acid methyl esters (FAMEs) were extracted with 200μL or 400μL n-hexane (RCI Labscan) and transferred into autosampler vials with inserts.

### GC-MS Analysis

GC-MS analysis was performed on an Agilent 7890B gas chromatograph equipped with an Agilent 5977A mass spectrometer, an Agilent 7693 autosampler, and an Agilent DB-23 30m × 0.25mm × 0.25μm GC column (#122– 2332; Agilent). 1μL of sample was injected in splitless mode. Carrier gas helium was set at a constant flow rate of 1.6 mL/min. The inlet temperature was 220°C. The column temperature was maintained at 100°C for 1 min, and then increased to 180°C at 10°C/min, then held at 180 °C for 5 min. Finally, temperature was raised to 220°C at 5 °C/min, and held at 220°C for 3 min. The retention time of each FAMEs were identified when compared to the peaks of FAME standards that purchased from Nu-Chek Prep or by using Supelco 37 Component FAME Mix. Mass spectra of unidentified peaks were matched against NIST14 library. The response factors (RFs) for each FAMEs relative to the internal standard were determined from the constructed calibration curve. Quantitation was achieved by relating the peak area of each FAME with the spiked internal standard.

## Acknowledgements

We thank John T. Crowl and Daniel Capek for their contributions in the *C. elegans* genetic screen, Shohei Mitani for providing mutant strain, Ben Zhong Tang for providing lipid droplet dyes, Justin Law for FACS, Sean McKinney and Jetty Lee for technical advice, the Molecular Biology core at the Stowers Institute for Medical Research for whole-genome sequencing, Tom Rapoport and Gunther Hollopeter for comments. This work was supported by the Stowers Institute for Medical Research, and grants 662013, 16101614 and HKUST12/CRF/13G from the Hong Kong Research Grants Council to H. Y. Mak, grants U1402225 and 31571388 from the National Natural Science Foundation of China to P. Liu, and a grant CAS16SC01 from the Croucher Foundation to H. Y. Mak and P. Liu.

## Author contributions

Z. Cao was responsible for Fig. 1 (B, E, I-J), 2 (B-L), 3, 4, S1D-K, S2, S3 (B-F), S4 (A-K). Y. Hao was responsible for Fig. 1 (C-D, F, H), Y.Y. Lee was responsible for Fig. 2L, 3E, S3B, P. Wang and B.H. Kang were responsible for Fig. 4 (E-H), Fig. S4 (D-K), X. Li and J. Qu were responsible for developing the SRS system and Fig. 1 (B, I), K. Xie and P. Liu were responsible for Fig. 1K, W.J. Lam was responsible for Fig. S4 (L-N), Y. Qiu was responsible for generating SEIP-1 expressing COS7 cell lines, G. Shui was responsible for Fig. S1 (B-C). Y. Hao was responsible for the initial characterization of anti-SEIP-1 antibodies, all transgenes, and preliminary experiments for Figs. 1, 2, 4, S1, and S2. H.Y. Mak and Y. Hao conducted genetic screens, genetic mapping and molecular cloning of *C. elegans* mutants. H.Y. Mak wrote the manuscript with inputs from Z. Cao.

## Supplemental Figure Legends

**Figure S1.**
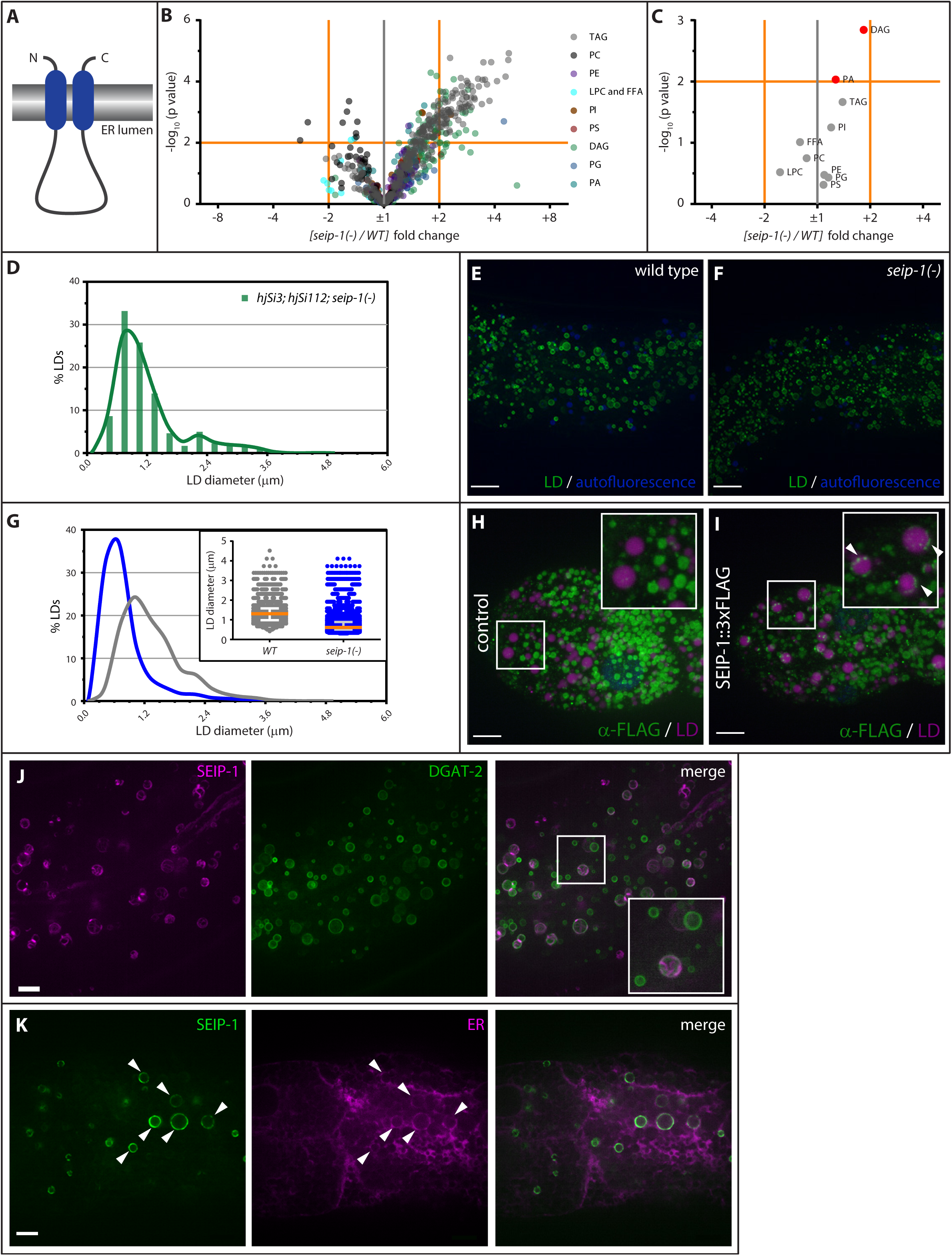
(A) Predicted membrane topology of SEIP-1. (B) Volcano plot of all 738 lipid species detected in WT or *seip-1(tm4221)* animals. (C) Volcano plot of total phospholipid (PC, PE, PI, PS, PG, PA and LPC), diacylglycerol (DAG), triacylglycerol (TAG) and free fatty acid (FFA) in wild-type (WT) or *seip-1(tm4221)* animals. (D) An example of the curve fitting model used in Fig. 1H, 3C and 4C. The curve was fitted twice using Fit Spline/LOWESS (20 points in smoothing window, 4000 segments) method in GraphPad Prism based on a histogram with a bin size of 0.3μm. (E) Visualization of LDs using the marker DHS-3::GFP (*hj120*, a knock-in allele at the endogenous *dhs-3* locus) in an otherwise wild type larval L4 stage animal. Autofluorescence from lysosome related organelles (LROs) is pseudocolored in blue. A projection of 4.5μm z stack centering at the second intestinal segment is shown. Scale bar = 10μm. (F) As in (E), but with a *seip-1(tm4221)* mutant animal. (G) Frequency distribution of LD diameter. The curve was fitted using the same method as in (D). Inset: a scatter plot showing the median and inter-quartile range of LD size. Total number of LDs measured: WT = 2,423; *seip-1(tm4221)* = 8,119. (H) A dissected wild type control animal was fixed and immunostained with mouse anti-FLAG antibodies and Alexa Fluor 488 anti-Mouse IgG. LDs were stained with LipidTox Red and pseudocolored magenta. Non-specific signals from the secondary antibodies are shown in green. Scale bar = 10μm. The boxed region was magnified 2x and shown in the inset. (I) As in (H), but with a seip-1::3xFLAG knock-in allele. Specific signals in the proximity of LDs are indicated by white arrowheads. (J) Visualization of SEIP-1::tagRFP (*hjSi434*) in a larval L4 stage animal. A projection of 3μm z stack centering at the second intestinal segment is shown. The LD marker GFP::DGAT-2 (*hjSi56*) is used. The tagRFP signal is pseudocolored magenta. The boxed region was magnified 2x and shown in the inset. Scale bar = 5μm. (K) Visualization of SEIP-1::GFP (*hjSi3*) in a larval L4 stage animal. A single focal plane centering at the second intestinal segment is shown. A luminal ER mCherry marker (*hjSi158*) is used and pseudocolored magenta. Scale bar = 5μm. The boxed area was magnified 2x and shown in the inset.

**Figure S2.**
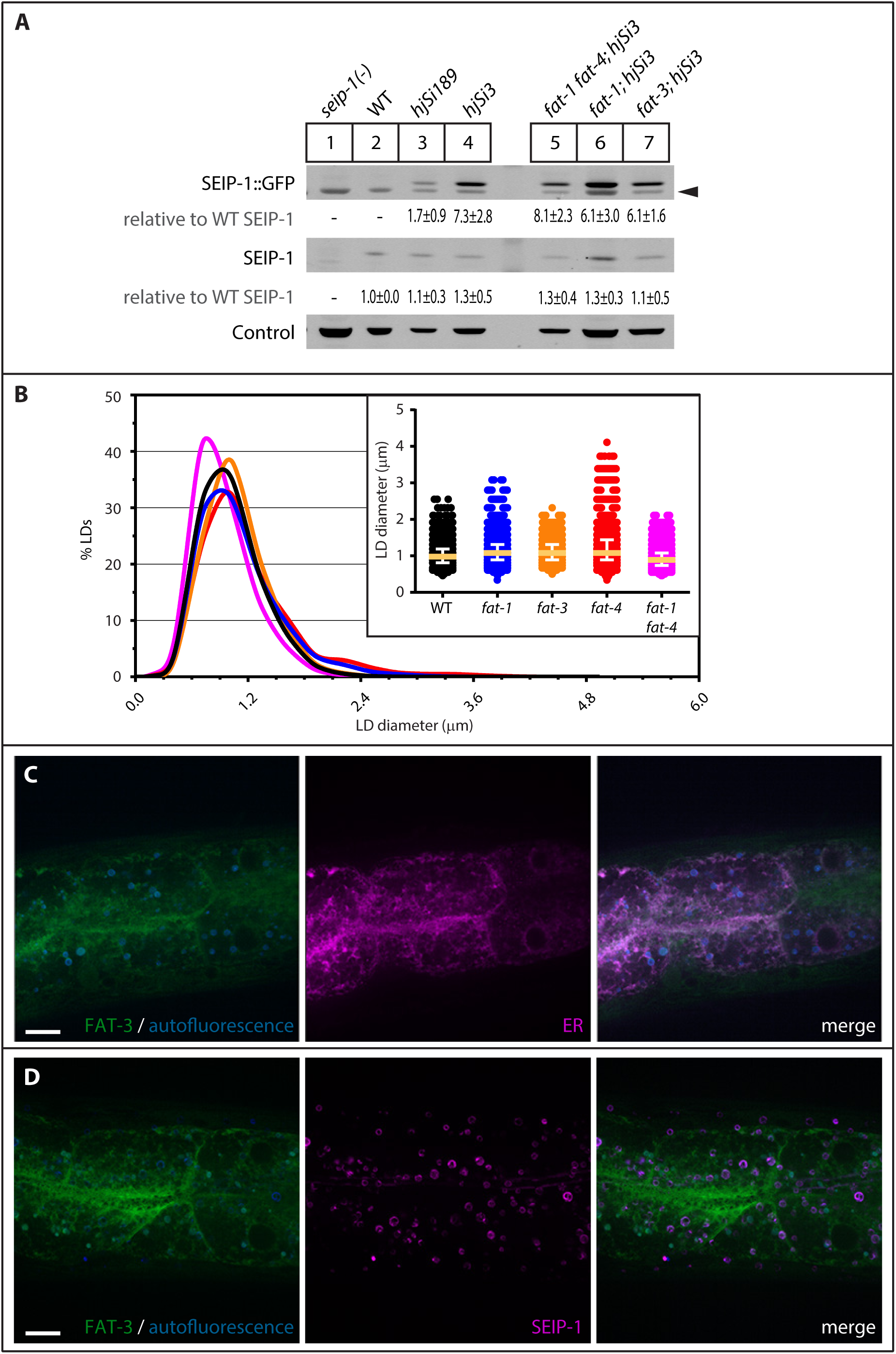
(A) The expression levels of SEIP-1::GFP and endogenous SEIP-1 were determined by SDS-PAGE and immunoblotting with anti-SEIP-1 antibodies. A non-specific band at ∼95kDa served as a loading control. The expression level of endogenous SEIP-1 in wild type (WT) animals served as a reference for normalization of signals in other samples. The mean ± SD from three independent experiments (including the blot in Figure 1B) is shown. The arrowhead indicates a non-specific band. (B) Frequency distribution of LD diameter. The curve was fitted using the same algorithm as in Fig. S1D. Inset: a scatter plot showing the median and inter-quartile range of LD size. Total number of LDs measured: WT = 4,600; *fat-1(wa9)* = 3,269; *fat-3(ok1126)* = 2,221; *fat-4(wa14)* = 2,577; *fat-1(wa9) fat-4(wa14)* = 3,293. (C) Visualization of FAT-3::GFP (*hjSi222*) in a larval L4 stage *fat-3(ok1126)* animal. The *hjSi222* transgene rescues the *fat-3* mutant phenotypes. A projection of 3μm z stack centering at the second intestinal segment is shown. Autofluorescence from lysosome related organelles is pseudocolored blue. The ER is marked by a tail-anchored mRuby red fluorescent protein that is expressed exclusively in the intestine (*hjSi127*). The mRuby signal is pseudocolored magenta. Scale bar = 10μm. (D) Visualization of both FAT-3::GFP (*hjSi222*) and SEIP-1::tagRFP (*hjSi434*) in a WT larval L4 stage animal. A projection of 2μm z stack centering at the second intestinal segment is shown. Autofluorescence from lysosome related organelles is pseudocolored blue. The tagRFP signal is pseudocolored magenta. Scale bar = 10μm.

**Figure S3.**
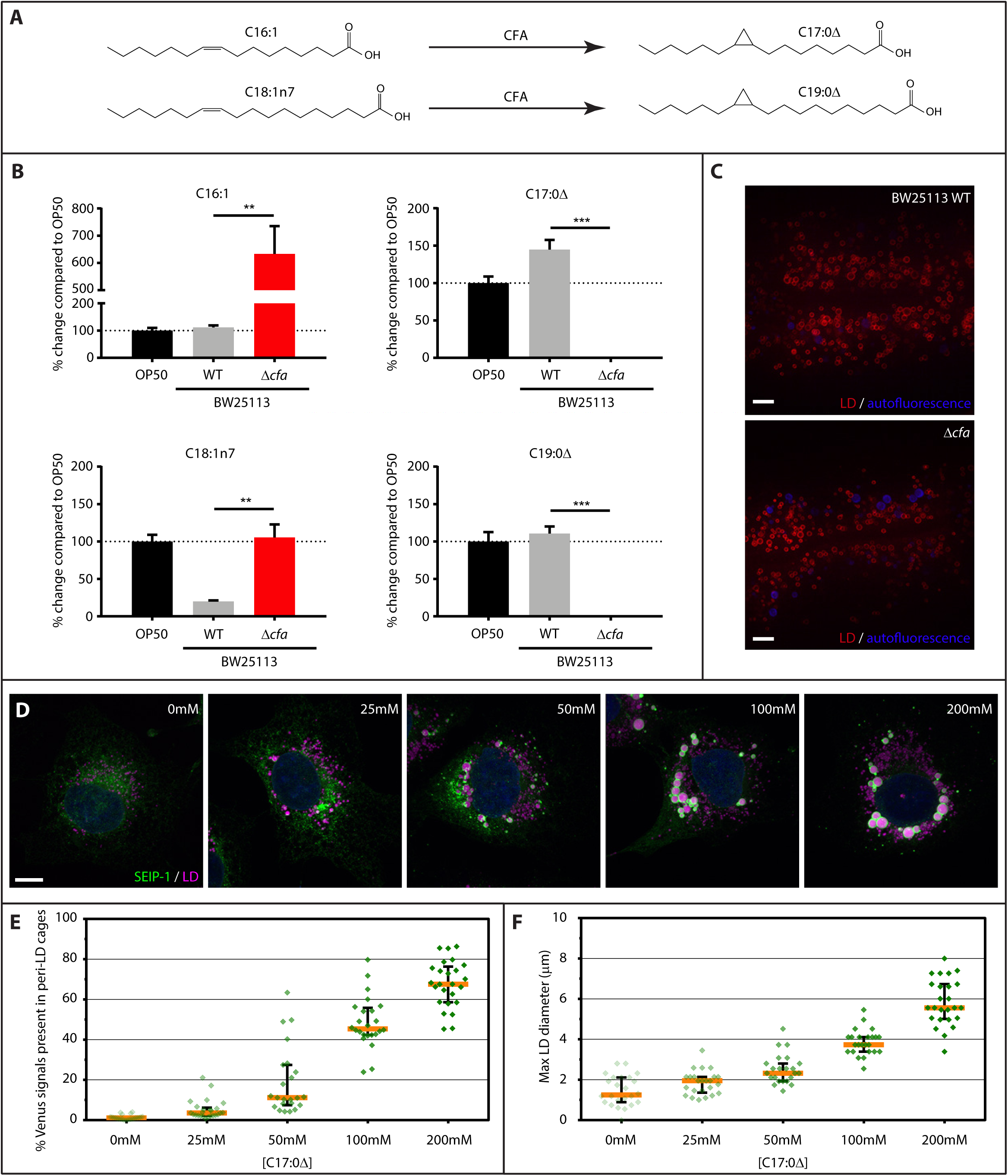
(A) The biosynthetic pathway for cyclopropane fatty acids in *Escherichia coli*. (B) The normalized abundance of cyclopropane fatty acids and their mono-unsaturated precursors in OP50, BW25113 WT and BW25113 *Δcfa E. coli*. Mean ± SEM of 3 independent samples is shown. **p < 0.01; ***p < 0.001 (unpaired t-test). (C) Visualization of LDs using the marker mRuby::DGAT-2 (*hjSi112*) in wild type larval L4 stage animals fed on BW25113 WT or *Δcfa E. coli*. Autofluorescence from lysosome related organelles (LROs) is pseudocolored in blue. A projection of 4.5μm z stack centering at the second intestinal segment is shown. Scale bar = 5μm. (D) COS7 cells stably expressing SEIP-1::Venus were supplemented with increasing concentration of cis-9,10-methylenehexadecanoic acid (17:0Δ). LDs were stained with LipidTox Red (pseudocolored magenta) and nuclei were stained by Hoechst 33342 (in blue). A projection of 4.5μm z stack is shown. Scale bar = 10μm. (E) The effects of FAs supplementation on the percentage of Venus signals in peri-LD cages. A scatter plot derived from at least 23 cells of each condition is shown. Median with interquartile range is displayed. (F) As in (E), but the effect of FA supplementation on the maximum LD diameter is shown.

**Figure S4.**
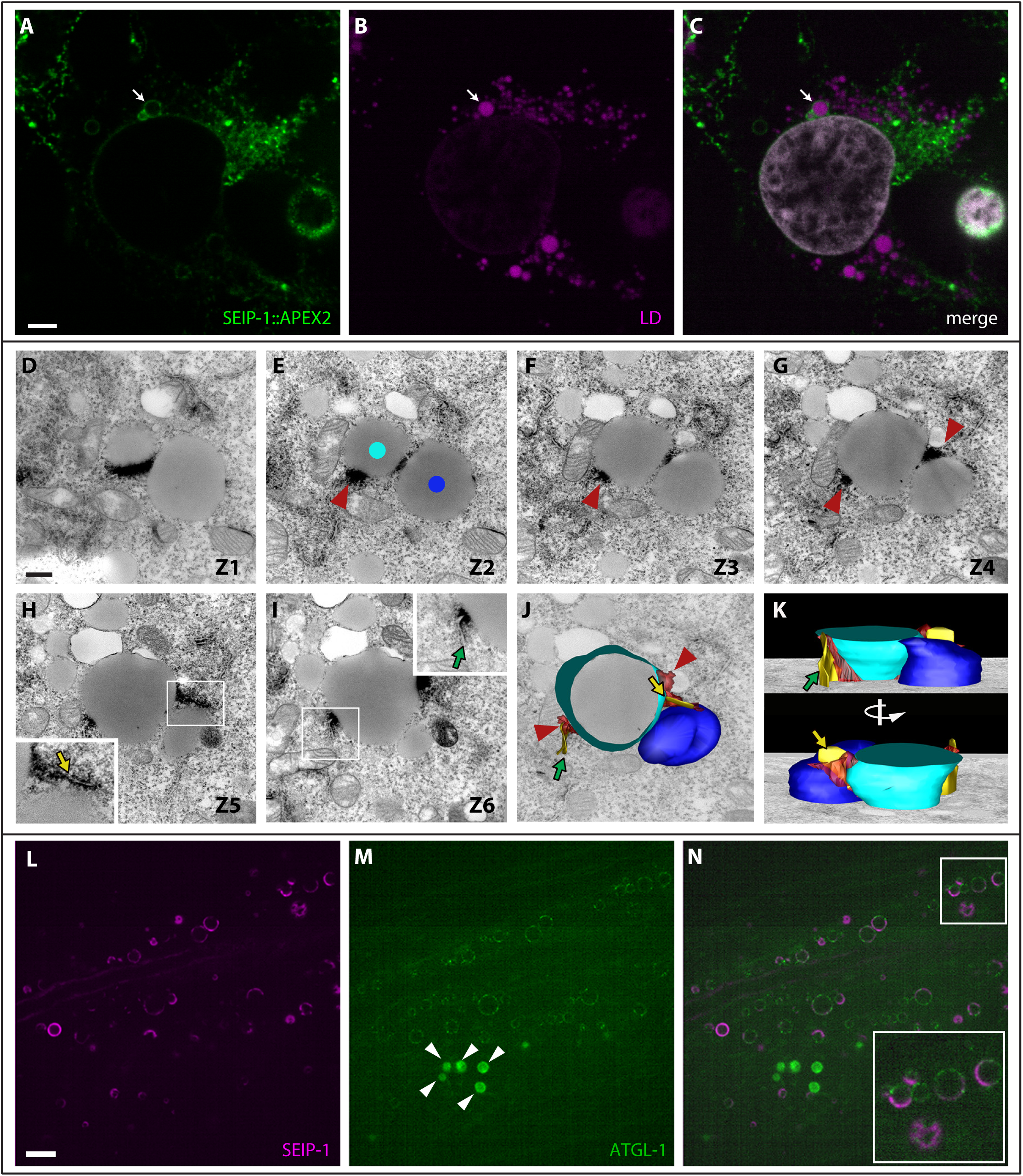
(A) COS7 cells stably expressing SEIP-1::V5-APEX2 as in Figure 4F were subjected to immunostaining with anti-V5 antibodies. The cells were incubated with oleic acid prior to fixation. The white arrow indicates an enlarged LD surrounded by SEIP-1. Scale bar = 5μm. (B) As in (A), but with LDs stained with the lipid dye FAS (pseudo-colored magenta). (C) As in (A), but with SEIP-1, LD and Hoechst 33342 (pseudo-colored grey) signals merged. (D-K) Transmission electron micrographs of six consecutive sections (150nm thick) of a COS7 cell stably expressing a SEIP-1::V5-APEX2 fusion protein. Dark deposits that decorate SEIP-1 positive ER membranes immediately adjacent to LDs are marked in (E), (F), and (G) with red arrowheads. The insets in (H) and (I) show 2x magnification of the boxed areas in which dark deposits decorate SEIP-1 positive ER membrane extensions distal to LDs (arrows). The micrographs were captured at 25,000x and the scale bar in (E) indicates 0.5μm. A three-dimensional model in (J) was generated from the serial sections shown in (D-I). Two LDs (cyan and blue dots in (E)), ER membrane extensions (yellow and green arrows in the insets of (H) and (I), respectively) and ER membranes proximal to LDs (red arrowheads) are shown. (K) Side views of the model in (J) is shown. The bottom panel shows the model after 180° rotation along the vertical axis. The ER does not appear as tubules in the reconstruction because of the thickness of the sections. (L) Visualization of SEIP-1::tagRFP (*hjSi434*) in a larval L4 stage WT animal. A single focal plane centered at the second intestinal segment is shown. The tagRFP signal is pseudocolored magenta. Scale bar = 5μm. (M) As in (L), but with ATGL-1::GFP shown. The GFP coding sequence was knocked into the endogenous *atgl-1* locus by CRISPR. The fluorescence intensity of GFP is below that of autofluorescence from lysosome related organelles (indicated by arrowheads). (N) As in (L), but with GFP and tagRFP signals merged. The boxed area was magnified 2x and shown in the inset.

## References

1. Fujimoto, T. & Parton, R. G. Not just fat: the structure and function of the lipid droplet. Cold Spring Harb. Perspect. Biol. 3, (2011).

2. Mak, H. Y. Lipid droplets as fat storage organelles in Caenorhabditis elegans: Thematic Review Series: Lipid Droplet Synthesis and Metabolism: from Yeast to Man. J. Lipid Res. 53, 28–33 (2012).

3. Walther, T. C. & Farese, R. V., Jr. Lipid droplets and cellular lipid metabolism. Annu. Rev. Biochem. 81, 687–714 (2012).

4. Krahmer, N. et al. Phosphatidylcholine Synthesis for Lipid Droplet Expansion Is Mediated by Localized Activation of CTP:Phosphocholine Cytidylyltransferase. Cell Metab. 14, 504–515 (2011).

5. Tauchi-Sato, K., Ozeki, S., Houjou, T., Taguchi, R. & Fujimoto, T. The surface of lipid droplets is a phospholipid monolayer with a unique Fatty Acid composition. J. Biol. Chem. 277, 44507–44512 (2002).

6. Penno, A., Hackenbroich, G. & Thiele, C. Phospholipids and lipid droplets. Biochim. Biophys. Acta 1831, 589–594 (2013).

7. Yang, L. et al. The proteomics of lipid droplets: structure, dynamics, and functions of the organelle conserved from bacteria to humans. J. Lipid Res. 53, 1245–1253 (2012).

8. Cermelli, S., Guo, Y., Gross, S. P. & Welte, M. A. The lipid-droplet proteome reveals that droplets are a protein-storage depot. Curr. Biol. CB 16, 1783–1795 (2006).

9. O’Byrne, S. M. & Blaner, W. S. Retinol and retinyl esters: biochemistry and physiology. J. Lipid Res. 54, 1731–1743 (2013).

10. Thiam, A. R. & Beller, M. The why, when and how of lipid droplet diversity. J. Cell Sci. 130, 315–324 (2017).

11. Thul, P. J. et al. Targeting of the Drosophila protein CG2254/Ldsdh1 to a subset of lipid droplets. J. Cell Sci. 130, 3141–3157 (2017).

12. Wilfling, F. et al. Triacylglycerol synthesis enzymes mediate lipid droplet growth by relocalizing from the ER to lipid droplets. Dev. Cell 24, 384–399 (2013).

13. Robenek, H. et al. Compartmentalization of proteins in lipid droplet biogenesis. Biochim. Biophys. Acta 1791, 408–418 (2009).

14. Blanchette-Mackie, E. J. et al. Perilipin is located on the surface layer of intracellular lipid droplets in adipocytes. J. Lipid Res. 36, 1211–1226 (1995).

15. Xu, N. et al. The FATP1-DGAT2 complex facilitates lipid droplet expansion at the ER-lipid droplet interface. J. Cell Biol. 198, 895–911 (2012).

16. Jacquier, N. et al. Lipid droplets are functionally connected to the endoplasmic reticulum in Saccharomyces cerevisiae. J. Cell Sci. 124, 2424–2437 (2011).

17. Coleman, R. A. & Lee, D. P. Enzymes of triacylglycerol synthesis and their regulation. Prog. Lipid Res. 43, 134–176 (2004).

18. Kuerschner, L., Moessinger, C. & Thiele, C. Imaging of lipid biosynthesis: how a neutral lipid enters lipid droplets. Traffic Cph. Den. 9, 338–352 (2008).

19. Klemm, R. W. et al. A conserved role for atlastin GTPases in regulating lipid droplet size. Cell Rep. 3, 1465–1475 (2013).

20. Hu, J., Prinz, W. A. & Rapoport, T. A. Weaving the web of ER tubules. Cell 147, 1226–1231 (2011).

21. Magré, J. et al. Identification of the gene altered in Berardinelli–Seip congenital lipodystrophy on chromosome 11q13. Nat. Genet. 28, 365–370 (2001).

22. Cartwright, B. R. & Goodman, J. M. Seipin: from human disease to molecular mechanism. J. Lipid Res. 53, 1042–1055 (2012).

23. Chen, W. et al. Berardinelli-Seip Congenital Lipodystrophy 2/Seipin Is a Cell-Autonomous Regulator of Lipolysis Essential for Adipocyte Differentiation. Mol. Cell. Biol. 32, 1099–1111 (2012).

24. Payne, V. A. et al. The human lipodystrophy gene BSCL2/seipin may be essential for normal adipocyte differentiation. Diabetes 57, 2055–2060 (2008).

25. Yang, W. et al. Seipin differentially regulates lipogenesis and adipogenesis through a conserved core sequence and an evolutionarily acquired C-terminus. Biochem. J. (2013). doi:10.1042/BJ20121870

26. Fei, W. et al. Molecular characterization of seipin and its mutants: implications for seipin in triacylglycerol synthesis. J. Lipid Res. 52, 2136–2147 (2011).

27. Cai, Y. et al. Arabidopsis SEIPIN Proteins Modulate Triacylglycerol Accumulation and Influence Lipid Droplet Proliferation. Plant Cell 27, 2616–2636 (2015).

28. Fei, W. et al. Fld1p, a functional homologue of human seipin, regulates the size of lipid droplets in yeast. J. Cell Biol. 180, 473–482 (2008).

29. Salo, V. T. et al. Seipin regulates ER-lipid droplet contacts and cargo delivery. EMBO J. 35, 2699–2716 (2016).

30. Szymanski, K. M. et al. The lipodystrophy protein seipin is found at endoplasmic reticulum lipid droplet junctions and is important for droplet morphology. Proc. Natl. Acad. Sci. U. S. A. 104, 20890–20895 (2007).

31. Fei, W. et al. A role for phosphatidic acid in the formation of ‘supersized’ lipid droplets. PLoS Genet. 7, e1002201 (2011).

32. Wang, H. et al. Seipin is required for converting nascent to mature lipid droplets. eLife 5, (2016).

33. van Meer, G., Voelker, D. R. & Feigenson, G. W. Membrane lipids: where they are and how they behave. Nat. Rev. Mol. Cell Biol. 9, 112–124 (2008).

34. Antonny, B., Vanni, S., Shindou, H. & Ferreira, T. From zero to six double bonds: phospholipid unsaturation and organelle function. Trends Cell Biol. 25, 427–436 (2015).

35. Yamashita, A. et al. Acyltransferases and transacylases that determine the fatty acid composition of glycerolipids and the metabolism of bioactive lipid mediators in mammalian cells and model organisms. Prog. Lipid Res. 53, 18–81 (2014).

36. Sledzinski, T. et al. Identification of cyclopropaneoctanoic acid 2-hexyl in human adipose tissue and serum. Lipids 48, 839–848 (2013).

37. Tsirigos, K. D., Peters, C., Shu, N., Käll, L. & Elofsson, A. The TOPCONS web server for consensus prediction of membrane protein topology and signal peptides. Nucleic Acids Res. 43, W401–407 (2015).

38. Lundin, C. et al. Membrane topology of the human seipin protein. FEBS Lett. 580, 2281–2284 (2006).

39. Li, X. et al. Integrated femtosecond stimulated Raman scattering and two-photon fluorescence imaging of subcellular lipid and vesicular structures. J. Biomed. Opt. 20, 110501 (2015).

40. Zhang, S. O. et al. Genetic and dietary regulation of lipid droplet expansion in Caenorhabditis elegans. Proc. Natl. Acad. Sci. U. S. A. 107, 4640–4645 (2010).

41. Na, H. et al. Identification of lipid droplet structure-like/resident proteins in Caenorhabditis elegans. Biochim. Biophys. Acta 1853, 2481–2491 (2015).

42. Wolinski, H., Kolb, D., Hermann, S., Koning, R. I. & Kohlwein, S. D. A role for seipin in lipid droplet dynamics and inheritance in yeast. J. Cell Sci. 124, 3894–3904 (2011).

43. Watts, J. L. & Browse, J. Genetic dissection of polyunsaturated fatty acid synthesis in Caenorhabditis elegans. Proc. Natl. Acad. Sci. U. S. A. 99, 5854–5859 (2002).

44. Grogan, D. W. & Cronan, J. E. Cloning and manipulation of the Escherichia coli cyclopropane fatty acid synthase gene: physiological aspects of enzyme overproduction. J. Bacteriol. 158, 286–295 (1984).

45. Chang, Y. Y. & Cronan, J. E. Membrane cyclopropane fatty acid content is a major factor in acid resistance of Escherichia coli. Mol. Microbiol. 33, 249–259 (1999).

46. Baba, T. et al. Construction of Escherichia coli K-12 in-frame, single-gene knockout mutants: the Keio collection. Mol. Syst. Biol. 2, 2006.0008 (2006).

47. Kowarz, E., Löscher, D. & Marschalek, R. Optimized Sleeping Beauty transposons rapidly generate stable transgenic cell lines. Biotechnol. J. 10, 647–653 (2015).

48. Lam, S. S. et al. Directed evolution of APEX2 for electron microscopy and proximity labeling. Nat. Methods 12, 51–54 (2015).

49. Lee, J. H. et al. Lipid droplet protein LID-1 mediates ATGL-1-dependent lipolysis during fasting in Caenorhabditis elegans. Mol. Cell. Biol. 34, 4165–4176 (2014).

50. Young, S. G. & Zechner, R. Biochemistry and pathophysiology of intravascular and intracellular lipolysis. Genes Dev. 27, 459–484 (2013).

51. Fischer, J. et al. The gene encoding adipose triglyceride lipase (PNPLA2) is mutated in neutral lipid storage disease with myopathy. Nat. Genet. 39, 28–30 (2007).

52. Schweiger, M., Lass, A., Zimmermann, R., Eichmann, T. O. & Zechner, R. Neutral lipid storage disease: genetic disorders caused by mutations in adipose triglyceride lipase/PNPLA2 or CGI-58/ABHD5. Am. J. Physiol. Endocrinol. Metab. 297, E289–296 (2009).

53. Han, S., Binns, D. D., Chang, Y.-F. & Goodman, J. M. Dissecting seipin function: the localized accumulation of phosphatidic acid at ER/LD junctions in the absence of seipin is suppressed by Sei1p(ΔNterm) only in combination with Ldb16p. BMC Cell Biol. 16, 29 (2015).

54. Wolinski, H. et al. Seipin is involved in the regulation of phosphatidic acid metabolism at a subdomain of the nuclear envelope in yeast. Biochim. Biophys. Acta 1851, 1450–1464 (2015).

55. Wang, C.-W., Miao, Y.-H. & Chang, Y.-S. Control of lipid droplet size in budding yeast requires the collaboration between Fld1 and Ldb16. J. Cell Sci. 127, 1214–1228 (2014).

56. Grippa, A. et al. The seipin complex Fld1/Ldb16 stabilizes ER-lipid droplet contact sites. J. Cell Biol. 211, 829–844 (2015).

57. Teixeira, V. et al. Regulation of lipid droplets by metabolically controlled Ldo isoforms. J. Cell Biol. 217, 127–138 (2018).

58. Eisenberg-Bord, M. et al. Identification of seipin-linked factors that act as determinants of a lipid droplet subpopulation. J. Cell Biol. 217, 269–282 (2018).

59. Niu, S.-L. et al. Reduced G protein-coupled signaling efficiency in retinal rod outer segments in response to n-3 fatty acid deficiency. J. Biol. Chem. 279, 31098–31104 (2004).

60. Mitchell, D. C., Niu, S. L. & Litman, B. J. Optimization of receptor-G protein coupling by bilayer lipid composition I: kinetics of rhodopsin-transducin binding. J. Biol. Chem. 276, 42801–42806 (2001).

61. Pinot, M. et al. Lipid cell biology. Polyunsaturated phospholipids facilitate membrane deformation and fission by endocytic proteins. Science 345, 693–697 (2014).

62. Vanni, S., Hirose, H., Barelli, H., Antonny, B. & Gautier, R. A sub-nanometre view of how membrane curvature and composition modulate lipid packing and protein recruitment. Nat. Commun. 5, 4916 (2014).

63. Sonnenburg, J. L. & Bäckhed, F. Diet-microbiota interactions as moderators of human metabolism. Nature 535, 56–64 (2016).

64. Lin, C.-C. J. & Wang, M. C. Microbial metabolites regulate host lipid metabolism through NR5A-Hedgehog signalling. Nat. Cell Biol. 19, 550–557 (2017).

65. Li, X., Jiang, M., Lam, J. W. Y., Tang, B. Z. & Qu, J. Y. Mitochondrial Imaging with Combined Fluorescence and Stimulated Raman Scattering Microscopy Using a Probe of the Aggregation-Induced Emission Characteristic. J. Am. Chem. Soc. 139, 17022–17030 (2017).

66. Ramachandran, P. V., Mutlu, A. S. & Wang, M. C. Label-free biomedical imaging of lipids by stimulated Raman scattering microscopy. Curr. Protoc. Mol. Biol. 109, 30.3.1–17 (2015).

67. Zhang, P. et al. Proteomic study and marker protein identification of Caenorhabditis elegans lipid droplets. Mol. Cell. Proteomics MCP 11, 317–328 (2012).

68. Kang, B.-H. Electron microscopy and high-pressure freezing of Arabidopsis. Methods Cell Biol. 96, 259–283 (2010).

69. Zhang, S. O. et al. Genetic and dietary regulation of lipid droplet expansion in Caenorhabditis elegans. Proc. Natl. Acad. Sci. 107, 4640–4645 (2010).

